# A unifying framework for synaptic organization on cortical dendrites

**DOI:** 10.1101/771907

**Authors:** Jan H. Kirchner, Julijana Gjorgjieva

## Abstract

Dendritic synaptic inputs are organized into functional clusters with remarkable subcellular precision at the micron level. This organization emerges during early postnatal development through patterned spontaneous activity and manifests both locally where nearby synapses are significantly correlated, and globally with distance to the soma. We propose a biophysically motivated synaptic plasticity model to dissect the mechanistic origins of this organization during development, and elucidate synaptic clustering of different stimulus features in the adult. Our model captures local clustering of orientation in ferret vs. receptive field overlap in mouse visual cortex based on the cortical magnification of visual space. Including a back-propagating action potential explains branch clustering heterogeneity in the ferret, and produces a global retinotopy gradient from soma to dendrite in the mouse. Therefore, our framework suggests that sub-cellular precision in connectivity can already be established in development, and unifies different aspects of synaptic organization across species and scales.

---

Neurons in the developing brain are already precisely connected before sensory organs mature. It is widely accepted that spontaneous activity refines circuit connectivity to mature levels at the scale of single neurons and networks. Recent studies have revealed that spontaneous activity can also establish fine-scale organization of individual synapses within the dendritic arborizations of single neurons^(1,2)^. One striking example of this organization is functional synaptic clustering: synapses onto dendrites of pyramidal neurons that receive correlated input or encode a common sensory feature are spatially grouped. Synaptic clustering has been observed across brain regions, developmental ages and diverse species from rodent to primate^(1–10)^ and has multiple functional benefits; it compartmentalizes the dendrites of single neurons, enables supra-linear integration of inputs and shapes memory formation^(11)^. However, the mechanistic origins of synaptic clustering dependent on spontaneous activity during early postnatal development, and its relation to functional organization in the adult, remain elusive.

Interestingly, some *in vivo* studies have reported lack of fine-scale synaptic organization^(12–16)^, while others reported clustering for different stimulus features in different species. For example, dendritic branches in the ferret visual cortex exhibit local clustering of orientation selectivity but do not exhibit global organization of inputs according to spatial location and receptive field properties^(5,7)^. In contrast, synaptic inputs in mouse visual cortex do not cluster locally by orientation, but only by receptive field overlap, and exhibit a global retinotopic organization along the proximal-distal axis^(6)^. We presently do not understand the factors that underlie these scale- and species-specific differences.

Previous modeling studies of synaptic clustering have focused on different aspects than stimulus feature tuning and its organization. Many implement activity-dependent plasticity operating on slow time scales necessary for long-term memory formation, arguing that clustered configurations encode memories more robustly and efficiently^(17,18)^, increase a cell’s computational capacity^(19)^ and can link multiple memories across extended periods ^(20)^. Spike-timing-dependent plasticity has also been shown to drive clustering – albeit of an unusual nature involving synaptic efficacies – with limited biological relevance^(21)^. Alternatively, clustering can also emerge as the Bayes-optimal solution to a classical conditioning task with unreliable synaptic transmission^(22)^, which lacks a mechanism for organizing correlated inputs from different axons. Therefore, what is missing is a single framework to explain activity-dependent synaptic clustering of different stimulus features and across different scales in line with recent experiments in development and adulthood.

Here we propose a computational framework to unify experimental findings and make novel predictions about the emergence of fine-scale synaptic organization. Motivated by recent findings in early postnatal development, we developed a biophysically inspired model of synaptic plasticity based on a push-pull mechanism of interacting neurotrophic factors required for clustering^(3,8)^. We demonstrate that our neurotrophin model can be reduced to a more general plasticity rule that leads to functional synaptic clustering when stimulated with spontaneous retinal waves in a realistic scenario of visual system development. Importantly, the same model captures clustering of different stimulus features in adult ferret vs. mouse visual cortex based on the spread of receptive field centers in visual space, a parameter that depends on the cortical magnification factor of visual space. We can further explain differences between local and global organization on the dendritic tree by introducing a back-propagating action potential that attenuates away from soma. In addition to unifying diverse experimental results, our framework suggests that activity and plasticity driving clustering are already present in early development to wire immature circuits with the remarkable sub-cellular precision observed in adulthood.

## Results

### And activity-dependent neurotrophin model exhibits distance-dependent synaptic competition

To achieve functional synaptic organization, we proposed a biologically plausible implementation of a developmental plasticity mechanism based on the influential Yin-Yang hypothesis of neurotrophin action^(8,23)^. Our model considers two types of signaling molecules that interact in a ‘push-pull’ manner: *immature pro-neurotrophins* lead to synaptic depression and apoptosis of axons and neurons, while *mature neurotrophins* promote synaptic potentiation and the survival of neurons. An important neurotrophin ubiquitous in development is brain-derived neurotrophic factor (BDNF), which induces synaptic potentiation by binding to the TrkB (tropomyosin receptor kinase B) receptor^(23)^. In contrast, its immature form, proBDNF, binds preferentially to the p75^NTR^ receptor and induces synaptic depression^(23)^. ProBDNF and BDNF are involved in sub-cellular synaptic organization and the emergence and maintenance of synaptic clustering in the developing hippocampus^(3,8)^. Interestingly, proBDNF is more prevalent than BDNF in early developmental stages^(24)^, which may imply dominance of depression over potentiation. However, the relative amounts of proBDNF and BDNF are controlled by the cleaving protease matrix metallopeptidase 9 (MMP9) – other proBDNF cleaving enzymes exist^(8,25)^ – which can convert proBDNF into BDNF in an activity-dependent manner allowing appropriately activated synapses to be potentiated^(3,8)^.

Inspired by these experimental results, we developed a computational *neurotrophin model* for the activity-dependent interaction dynamics between proBDNF, BDNF and MMP9 during postnatal development (Fig. 1A and Methods). The postsynaptic release of proBDNF and BDNF into extracellular space depends on the opening of postsynaptic N-methyl-Daspartate (NMDA) channels and the subsequent influx of calcium^(8)^, which is known to spread postsynaptically upon synaptic activation in the developing brain^(26)^. Therefore, through the lateral spread of calcium, the activated synapse can exert a direct effect on a different nearby synapse by triggering neurotrophin release independent of presynaptic stimulation of that synapse^(25)^. MMP9 release is similarly coupled to neural activity^(25)^, but instead of spreading postsynaptically, it is co-localized with AMPA and NMDA receptors in excitatory synapses^(27)^. To capture this activity-dependent interaction of BDNF and proBDNF and their role in modulating the efficacy of synaptic inputs on local portions of the dendritic tree, we modeled MMP9 and calcium as local synapse-specific pre- and postsynaptic accumulators of neural activity, respectively (Fig. 1B and Methods).

**Figure 1:**
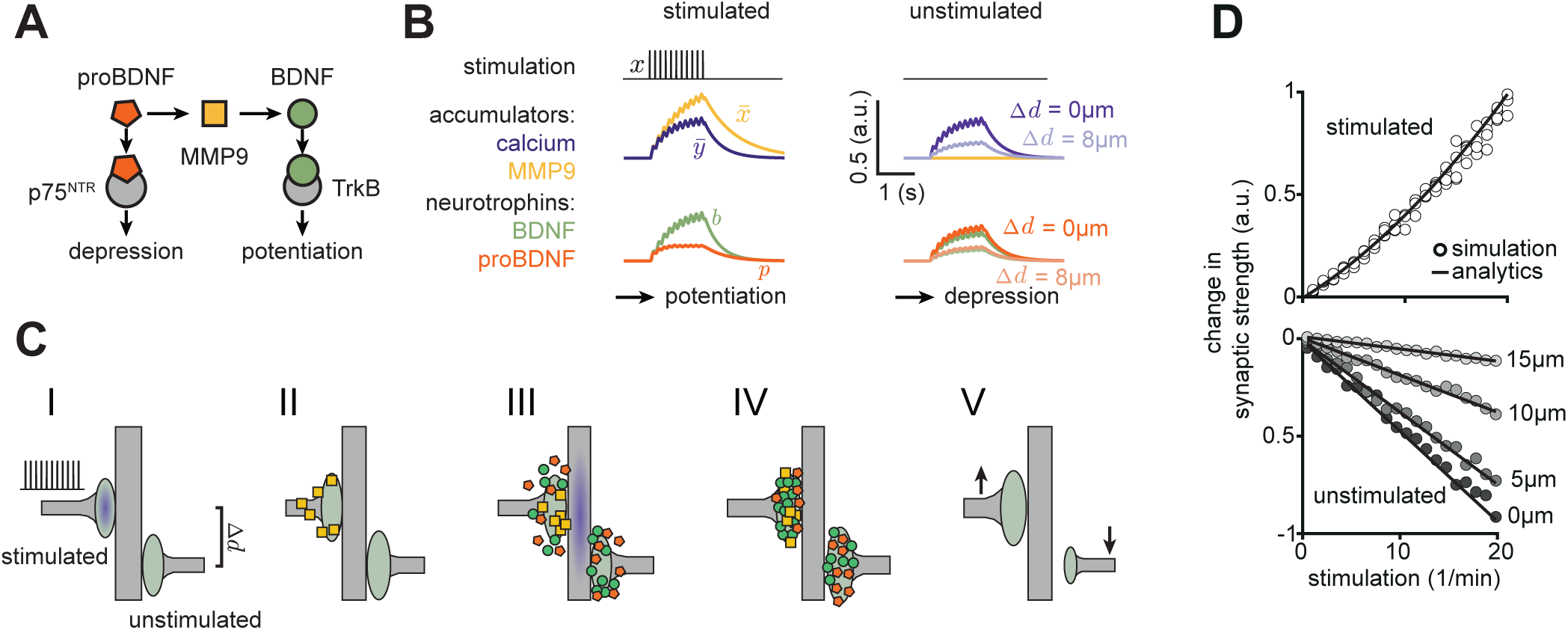
Distance-dependent synaptic competition in the neurotrophin model. (**A**) Model schematic: interactions between neu-rotrophins (BDNF and proBDNF), cleaving protease (MMP9) and neurotrophin receptors (TrkB and P75^NTR^). (**B**) Variables in **A** upon stimulation of one synapse only; the unstimulated synapse is positioned a distance Δ*d* µm away. (**C**) Outcome of synaptic stimulation: I. The left synapse is stimulated with a burst of action potentials. II. Presynaptic activation causes the local release of MMP9. III. Signal transduction into the postsynapse results in the spatially extended influx of calcium (purple shading). Calcium triggers the exocytosis of proBDNF and BDNF into extracellular space. IV. MMP9 differentially cleaves proBDNF into BDNF at the stimulated synapse. V. Repeating this pattern of unilateral stimulation potentiates the stimulated synapse, and depresses the unstimulated synapse. Symbols as in **A**. (**D**) Change in synaptic efficacy at the stimulated synapse (top) and the unstimulated synapse (bottom) as a function of input rate (in bursts per minute) and inter-synapse distance Δ*d*. The analytic solution follows from a linearized version of the model (see Methods).

We next sought to investigate the functional implications of our proposed neurotrophin model for the plasticity of interacting nearby synapses in a developmentally relevant scenario. Since during development bursts rather than single spikes are the units of information transmission^(3,28–30)^, we implemented a rate stimulation protocol inspired by the developing neuromuscular junction^(25)^ in which one synapse receives repeated 50-millisecond-long, continuous bursts of activation, while a second nearby synapse does not receive any input (Fig. 1B,C). In the neurotrophin model, MMP9 released at the stimulated synapse shifts the balance between proBDNF and BDNF in favor of BDNF (Fig. 1B; Table 1). Thus, the stimulated synapse potentiates, while the unstimulated synapse depresses as it remains dominated by proBDNF due to the absence of MMP9. This relationship, in which stimulated synapses potentiate and unstimulated synapses depress, is preserved for different stimulation rates (Fig. 1D) and is consistent with several experiments^(25,31,32)^. The resulting synaptic depression is attenuated with increasing inter-synaptic distance by a factor dependent on the spatial calcium spread constant (Fig. 1D and Methods). Therefore, consistent with experimental data^(3,33,34)^, our neurotrophin model provides a mechanistic explanation for the activity-dependent competition between differentially stimulated synapses as a function of inter-synaptic distance.

**Table 1:** Parameters of the proposed model along with nominal values used for simulations, unless stated otherwise. Citations indicate a free parameter fitted to experimental data. * indicates a parameter that is derived from the other parameters.

| Parameter | Variable | Value |
| --- | --- | --- |
| Synaptic efficacy time constant<sup>(74)</sup> | $\tau_W$ | 6 s |
| proBDNF and BDNF time constants<sup>(32)</sup> | $\tau_P, \tau_B$ | 5 ms |
| Postsynaptic calcium time constant<sup>(26)</sup> | $\tau_Y, \tau_u$ | 300 ms |
| Extracellular cleaving time constant | $\tau_M, \tau_v$ | 600 ms |
| Constitutive percent of BDNF of amount of total neurotrophins released<sup>(73)</sup> | $\eta$ | 45% |
| MMP9 efficiency constant<sup>(25)</sup> | $\phi$ | 3 |
| Heterosynaptic offset* | $\rho$ | $\rho = \frac{2\eta-1}{2(1-\eta)}$ |
| Minimal model synaptic efficacy time constant* | $\tau_w$ | $\tau_w = \tau_W \frac{1}{2(1-\eta)}$ |
| Standard deviation of calcium spread<sup>(26)</sup> | $\sigma_c$ | 6 $\mu\text{m}$ |
| Density of synapses<sup>(61,75)</sup> | $\nu$ | 0.2 $\mu\text{m}^{-1}$ |
| Spread of receptive field centers for ferret and mouse<sup>(6,7)</sup>, in visual space | $\sigma_p$ | 5.3°, 26° |
| Semi-minor and semi-major axis of Gabor receptive field<sup>(6,7)</sup> | $\sigma_{min}, \sigma_{maj}$ | 1°, 2° |
| LN model parameters<sup>(3,7)</sup> | $a, b$ | 0.2 Hz, 9.4 |
| Turnover threshold below which a synapse is replaced | $W_{thr}$ | 0.02 |
| Threshold for bAP generation<sup>(52)</sup> | $B_{thr}$ | 25 |
| Unattenuated amplitude of bAP<sup>(51,52)</sup> | $B_{amp}$ | 5 |

### The neurotrophin model exhibits burst-timingdependent plasticity

We next asked whether our neurotrophin model can be linked to correlated-based Hebbian learning rules operating in development. According to these rules, input correlations affect the sign and magnitude of synaptic plasticity, with asynchronous inputs leading to synaptic depression^(3)^. One such phenomenological plasticity rule – termed Burst-Timing-Dependent-Plasticity, or BTDP – characterized at the cellular level in the developing visual system finds that synchronous pre- and postsynaptic bursts induce synaptic potentiation, while pre- and postsynaptic bursts separated by more than a second induce synaptic depression^(35)^. BTDP differs from Spike-Timing-Dependent-Plasticity (STDP)^(36)^ in two major ways: first, the rule is symmetric so that it does not matter whether the pre- or the postsynaptic signal comes first. Second, BTDP operates on much longer timescales than STDP. We relate the change in synaptic efficacy in our neurotrophin model to BTDP under the aforementioned burst-stimulationprotocol^(35)^ (Fig. 2A). We find that temporal offsets below one second in our model yield synaptic potentiation due to the relative dominance of BDNF-induced signaling over proBDNF-induced signaling, while longer offsets lead to depression (Fig. 2B-D).

**Figure 2:**
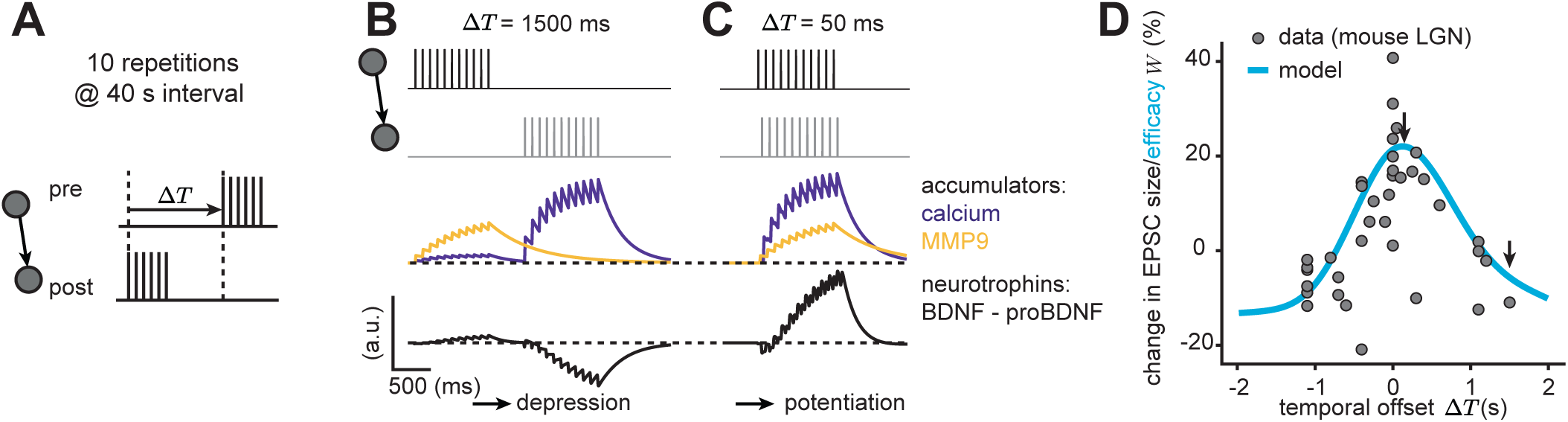
The neurotrophin model is consistent with burst-timing-dependent plasticity. (**A**) A burst-timing dependent stimulation protocol where the timing between pre- and postsynaptic bursts is varied^(35)^. (**B,C**) Accumulator and neurotrophin variables (Fig. 1B) under the burst pairing protocol with Δ*T* = 1.5s (**B**) and Δ*T* = 0.05 s (**C**). (**D**) Percentage change in synaptic efficacy as a function of burst offset, Δ*T* (data reproduced from ref. ^(35)^). The arrows mark the temporal offsets displayed in **B,C**.

The surprising link to BTDP enabled us to mathematically reduce our neurotrophin model to a Hebbian learning rule where synaptic change depends on accumulated presynaptic (‘pre’) and postsynaptic (‘post’) activity and a constant ‘offset’ related to the initial ratio of proBDNF to BDNF in the absence of extracellular conversion (see Methods),

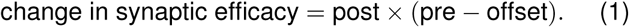

This reduced model has the advantage over the full neurotrophin model that it can be analyzed mathematically (see Methods), but is still completely consistent with the BTDP rule as well as distance-dependent synaptic competition (Supplementary Fig. 8). Since this reduced model makes a minimal set of assumptions for the correlation- and distance-dependent plasticity of individual dendritic synapses, we refer to it as a *minimal model* of local synaptic plasticity and use it for the remainder of this study. Moreover, this general form has the advantage that it can be implemented by other interacting molecules than the developmentally relevant proBDNF and BDNF (see Discussion).

### Input correlations and synaptic density determine synaptic competition

We next tested how our minimal model modulates the efficacy of individual synapses based on local interactions between them by systematically varying input correlation and synaptic density on the dendritic branch (Fig. 3A). For randomly distributed synapses receiving Poisson input trains with homogeneous pairwise correlations, we derived a mathematical relationship that describes the change in synaptic efficacy as a function of input correlation and synaptic density (Fig. 3B). We identified two regimes: (i) all synapses potentiate – if input correlation is high and independent of synaptic density (Fig. 3C, diamond), or if input correlation and synaptic density are both low (Fig. 3C, star), and (ii) synapses compete resulting in the selective potentiation of only a subset of synapses – if input correlation is low but synaptic density is high (Fig. 3C, triangle). The minimal model (Eq. 1) thus resembles the Hebbian covariance rule^(37)^, but with the additional feature that synaptic efficacy is modulated by the local density of synaptic inputs on the dendrite. Our results also hold for more realistic heterogeneous correlations across pairs of synapses, selected from a biologically realistic skewed distribution^(38)^ (Fig. 3D and Methods).

**Figure 3:**
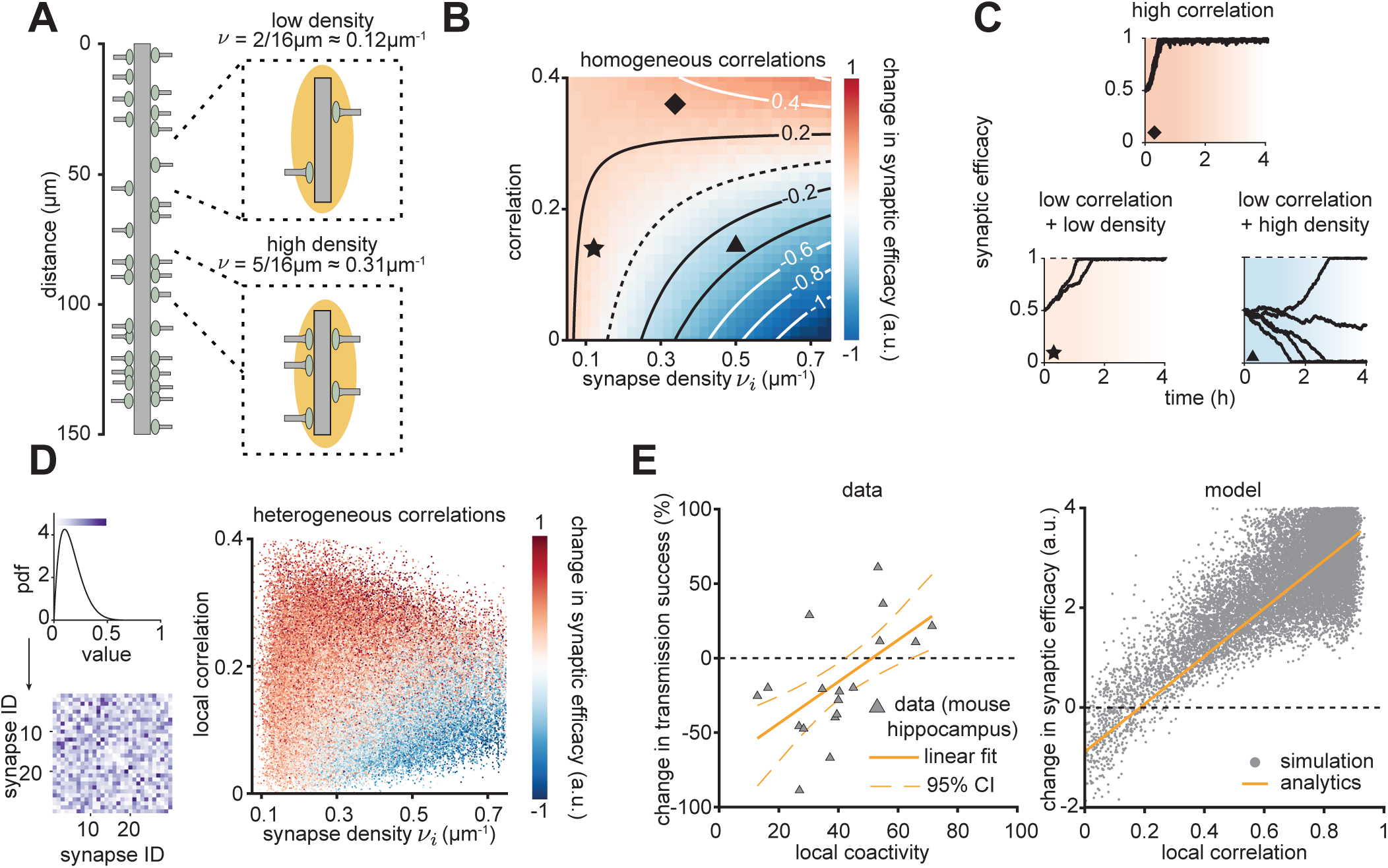
Relating synaptic change to synaptic density and input correlation. (**A**) Schematic with randomly distributed synapses along a linear dendritic branch. All synapses receive Poisson input trains with identical pairwise correlations (homogeneous, **B,C**) or from a distribution (heterogeneous, **D,E**). Inset: low (top) and high (bottom) density of synapses, *ν*. (**B**) Change in synaptic efficacy after 12 minutes of simulation as a function of input correlation and density. Contour lines are derived analytically while colors are determined by simulation. Color indicates negative (blue) or positive (red) change in synaptic efficacy averaged over time and synapses. (**C**) Synaptic efficacy over time for three different combinations of density and correlation (color and symbols as in **B**). (**D**) Left: A skewed pairwise input correlation distribution (top) and an example correlation matrix (bottom). Right: Change in synaptic efficacy after 24 minutes of simulation as a function of synaptic density and the correlation averaged for nearby (< 6 µm) synapses. (**E**) Left: Experimental data from developing mouse hippocampus slices of the percentage change in transmission success as a function of local coactivity (related to input correlation), defined as the percent of events that occurred within one second of nearby events (gray triangles, reproduced from ref. ^(3)^). Right: Change in synaptic efficacy after 12 minutes of simulation as a function of the heterogeneous correlation averaged for nearby (< 6 µm) synapses in the simulations. Gray dots indicate individual synapses at a random position on the dendrite and random initial efficacy. Orange lines denote the analytical predictions for homogeneous correlations (see Methods).

The synaptic competition that our model produces for low input correlations and high synaptic density (Fig. 3B,C, triangle) resembles a local *out-of-sync-lose-your-link* plasticity rule characterized experimentally in the developing visual cortex and hippocampus^(3)^. According to this rule, synaptic inputs on developing pyramidal dendrites either increase or decrease the success rate of synaptic transmission depending on their synchronization with their neighbors (Fig. 3E, left). For the same synaptic density corresponding to the developmental stage of the animal (0.3 μm^*-*1^) as in the original experiments^(3)^, our plasticity model generates a similar monotonically increasing relationship between the average correlation of nearby synapses and the change in synaptic efficacy, which we use as a measure of synaptic transmission success (Fig. 3E, right). Therefore, our model captures the depression at weakly correlated inputs embodying out-of-sync-lose-your-link plasticity.

### Retinal wave input and synaptic turnover drive synaptic orientation clustering

Next we sought to determine the potential of our minimal model to drive synaptic organization under realistic conditions in the developing visual cortex, which exhibits patterns of spontaneous activity propagated from the sensory periphery^(39,40)^. Therefore, we used simulated retinal waves generated from a published model^(41)^ as inputs. To convert retinal waves to cortical synaptic input, we implemented a two-stage linear-nonlinear (LN) model^(42)^ (Fig. 4A). The LN model has two components, a filter that linearly filters the retinal waves and a nonlinearity that produces spiking probability as a function of the filtered retinal wave input. For the linear filter, we use a spatial Gabor filter since recent studies have found that the receptive fields of spines on pyramidal neurons in the adult mouse visual cortex are oriented with positive and negative elongated subregions^(6)^. Individual pyramidal neurons in the mouse and ferret visual cortex are already orientation selective at eye opening^(43–45)^. Therefore, we assumed that individual synaptic inputs can already be described by Gabor filters in early development (though we also explored the formation of Gabors; Supplementary Fig. 9) and investigated whether our model can use patterns of spontaneous activity to organize them on the dendritic tree. Each Gabor filter was characterized by two parameters: the receptive field center, described by polar coordinates, and its orientation, with a value between 0^°^ and 360^°^ (Fig. 4A).

**Figure 4:**
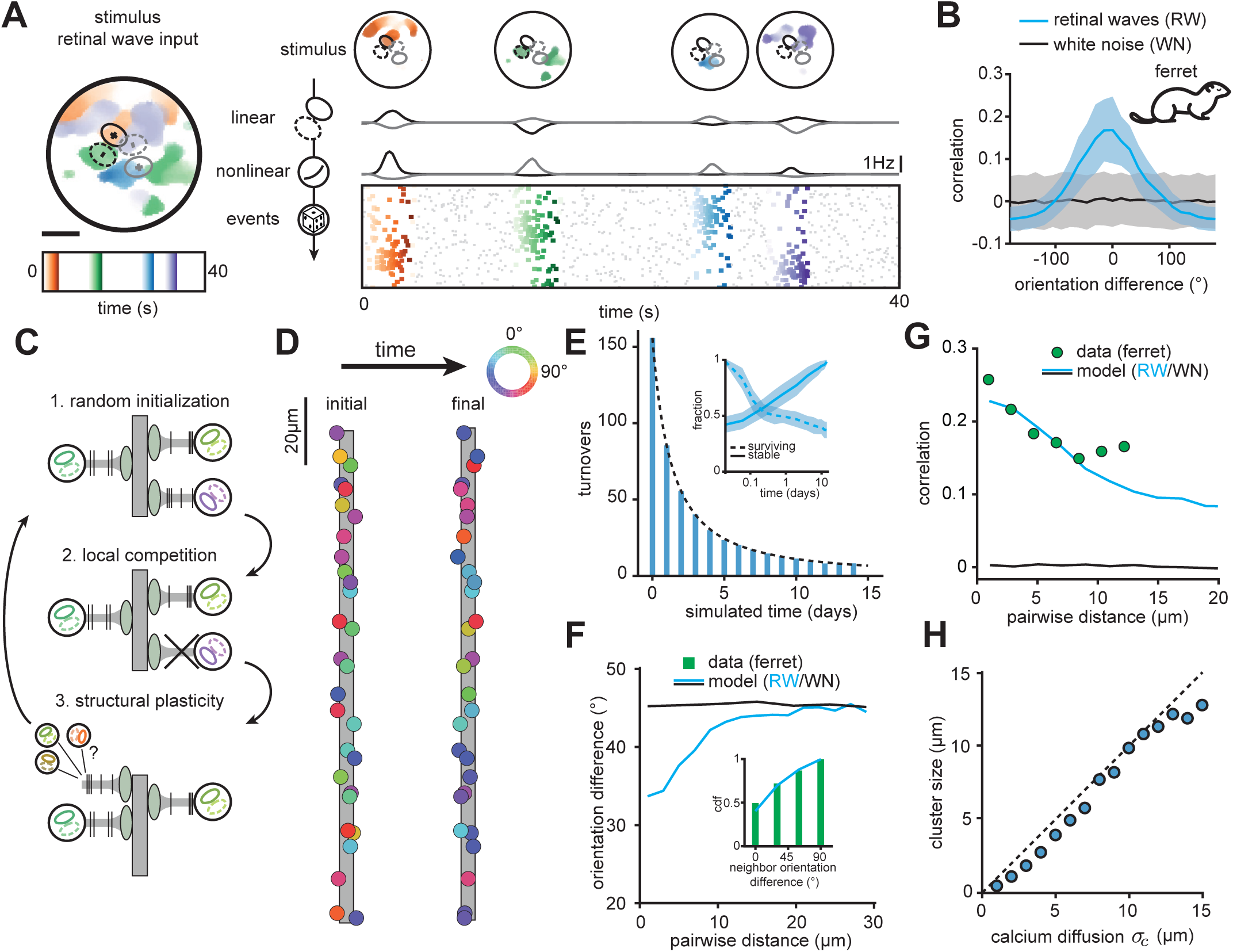
Retinal wave input and synaptic turnover generate orientation clustering on an L2/3 dendrite in a model of ferret V1. (Left) Illustration of forty seconds of retinal wave input. Different color gradients indicate different time points, scale bar is 30° in visual space. Superimposed are two schematics of Gabor receptive field filters with positive and negative components. (Right, from top) Stimulus (four spontaneous retinal waves), linearly filtered response, nonlinear output and Poisson events for the two oriented Gabors on the left. Rows correspond to different receptive fields, sorted from top to bottom by orientation. Gray events represent background noise. (**B**) Correlations as a function of the difference in orientation of receptive fields stimulated with retinal waves and white noise. Shaded area indicates standard deviation. (**C**) Local correlation- and distance-dependent competition together with structural plasticity mechanism drive the formation of synaptic clusters. (**D**) Example of the emergence of orientation clustering on a linear dendrite with retinal waves over two simulated weeks. (**E**) Number of turnovers as a function of simulated days. Inset: Semi-log plot of the survival fraction (synapses present at the beginning of simulation) and the stable fraction (synapses present at the end of the simulation) as a function of simulated days. Shaded area indicates standard deviation. (**F**) Orientation difference between pairs of synapses as a function of distance in the model with retinal wave input or white noise input. Inset: Cumulative fraction of orientation difference between an individual synapse and its nearby neighbors (less than 3 µm away) in the model and experiments in the ferret visual cortex (reproduced from ref. ^(5)^). Note that we compute the difference after taking the orientation modulo 180° (see Methods). (**G**) Correlations between pairs of synaptic inputs driven in the model by retinal waves or white noise and correlations between calcium signals of spontaneous synaptic activity in the ferret visual cortex (reproduced from ref. ^(7)^) as a function of synaptic distance. (**H**) Cluster size computed as the standard deviation of the best Gaussian fit to the pairwise correlation vs. distance (see Methods and **G**), as a function of the postsynaptic calcium spread constant *σ*_*c*_. Dashed line indicates the identity line. For the simulations in this figure, receptive field centers in visual space were distributed as in the ferret visual cortex^(7)^.

As the second stage of the LN model, we used an exponential nonlinearity to convert the linearly filtered retinal waves into instantaneous firing rates from which activity is generated via a Poisson process (see Methods). Thus, in the LN model, a synapse probabilistically receives bursts of action potentials when the Gabor filter is appropriately stimulated by a retinal wave traveling in the direction that matches the filter orientation (Fig. 4A). Since activation of a synapse depends on the appropriate stimulation of its associated receptive field by retinal waves, synapses whose receptive fields have nearby centers in visual space (as is the case in the ferret visual cortex^(7)^) and a small difference in orientation receive correlated input (Fig. 4B). White noise stimulation does not produce correlations for any orientation difference due to the lack of spatiotemporal structure to consistently activate a receptive field (Fig. 4B).

Next, we investigated the functional implications of correlation- and distance-dependent plasticity in our minimal model on synaptic organization with retinal wave input. We distributed synaptic inputs with randomly oriented receptive fields and centers distributed in visual space as measured experimentally^(7)^ on a non-branching, linear dendrite and stimulated them with action potentials after passing retinal waves through an LN model (Fig. 4D, left). Neighboring synapses with mismatched orientations receive uncorrelated input as they are rarely activated by the same retinal wave and will consequently depress. To prevent their irrevocable elimination, we further introduced an activity-dependent mechanism of structural plasticity, which preserves the total number of synaptic inputs on a dendritic branch^(46,47)^ (Fig. 4C). Upon the removal of a synapse, a new synapse was placed at a random position on the dendritic branch having a receptive field with random orientation and visual space location sampled from the experimentally characterized distribution^(7)^ (see Methods).

Throughout the simulation, synapses compete, and either stabilize or depress and turn over until nearby synapses share a similarly oriented receptive field. With ongoing simulation, the number of turnovers per day decreases rapidly with all remaining turnovers due to a small fraction of synapses (Fig. 4E). Despite a massive turnover during the first three days of the simulation, approximately 40% of synapses that are present at the beginning of the simulation do not experience any turnover and form a scaffold around which the remaining synapses stabilize (Fig. 4E, inset).

We find that nearby synapses in the stable state share similarly oriented receptive fields, a type of clustering that we call *orientation clustering* and which does not emerge with white noise stimulation (Fig. 4D, right and F). Orientation clustering has recently been reported in layer 2/3 of the adult ferret visual cortex recorded *in vivo*^(5,7)^. In addition, the correlations between pairs of synapses trained with retinal wave input (but not white noise) in our simulations decay with distance at the same rate as measured from spontaneous activity in experiments (Fig. 4G). When using this relationship to characterize the size of the formed clusters, we find that cluster size strongly depends on the spatial spread constant of postsynaptic calcium, with bigger clusters produced by broader calcium spread (Fig. 4H, Supplementary Fig. 10). Therefore, we predict that the different sizes of orientation clusters found in different species (ferrets^(7)^ or macaques^(10)^) and the variability in different cells in the same animal^(7)^ could be the result of different amounts of postsynaptic calcium spread.

In summary, our model implements an iterative process that, starting from synaptic inputs with random locations in visual space and random orientations, converges to a stable configuration where similarly oriented and functionally correlated inputs are locally clustered on a dendritic branch. This process occurs through the selective elimination of synaptic inputs that have mismatched orientation preference compared to their neighbors. The key factors driving this process are (1) distance-dependent competition, (2) dependence on correlated input, which is determined by realistic inputs during development in the form of retinal waves, and (3) synaptic turnover.

### Clustering of different features in mouse and ferret

Synapses are not clustered for every stimulus feature^(12,13,16)^. In stark contrast to the ferret or the macaque, nearby synapses on pyramidal neuron dendrites in the mature mouse visual cortex do not share a preference for the same orientation^(6,14,15,49)^. However, despite this lack of orientation clustering, the mouse visual cortex exhibits correlated activity in nearby synapses that emerges during development^(3)^. We propose that this functional organization could be the result of synaptic clustering for receptive field overlap, rather than orientation. Such clustering of synapses with overlapping receptive fields has been demonstrated recently in experiments in adult mouse visual cortex^(6)^. We refer to this type of clustering as *overlap clustering* to contrast it with the orientation clustering observed in the ferret.

What determines whether synapses exhibit orientation or overlap clustering? One striking difference between the two species is that cortical magnification of visual space in the ferret visual cortex is one order of magnitude larger than in the mouse^(48,50)^. As a result, a pyramidal neuron in the ferret visual cortex receives inputs from a considerably smaller region of visual space, which would promote orientation clustering (Fig. 5A, top). A pyramidal neuron in the mouse visual cortex, however, receives input from a broader region of visual space, thus favoring more intricate arrangements of receptive fields that would still produce correlated synaptic activity even without orientation clustering (Fig. 5A, middle). These differences can be captured by the relative broadness of the distribution of receptive field centers characterized experimentally in the two species^(6,7)^, a parameter that we call *receptive field center spread* (Fig. 5A, bottom).

Indeed, using a larger spread of receptive field centers measured in the mouse visual cortex^(6)^ leads to overlap clustering after training with retinal waves, quantified by the decreasing overlap between the receptive fields of pairs of synapses as a function of inter-synaptic distance (Fig. 5B,C and Methods). We observe an even stronger overlap between nearby synapses than experimentally reported because the model receptive fields are more regular than the experimentally estimated receptive fields. Adding irregularities to the Gabor synaptic filters improves the match between model and data (Supplementary Fig. 11). Overlap clustering also leads to functional clustering as observed in development^(3)^ (Fig. 5D). The resulting synaptic input configuration is stable (Supplementary Fig. 12) and, in agreement with experimental data, lacks orientation clustering (Fig. 5B,E).

**Figure 5:**
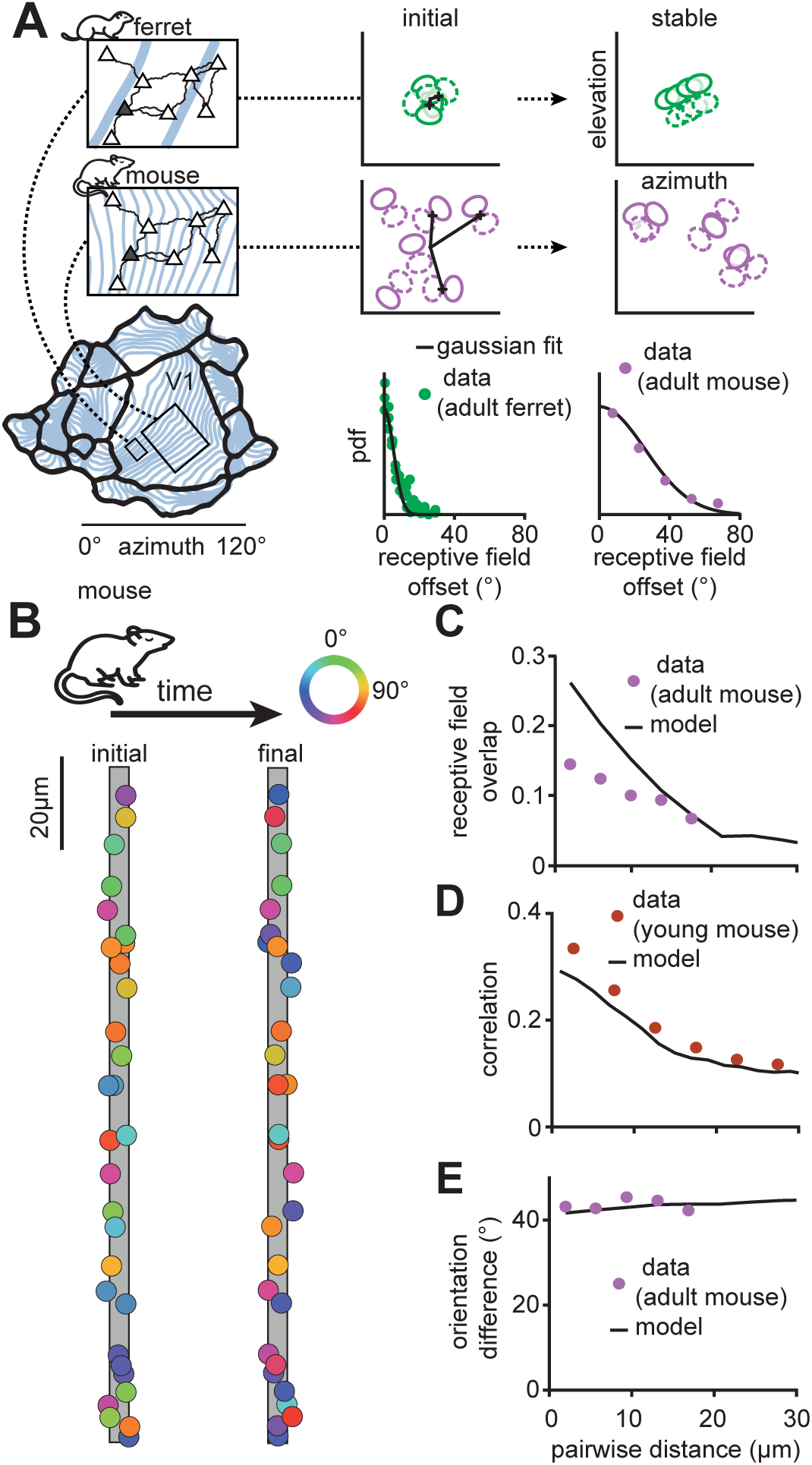
Overlap clustering, but not orientation clustering, emerges on an L2/3 dendrite in a model of mouse V1. (**A**) Schematic of the anatomical argument for two qualitatively different types of clustering in mouse and ferret. 1 mm of visual cortex spans a larger region of the total visual space (bottom left, blue lines represent iso-contours at ∼ 7°^(48)^) in the mouse than in the ferret. Since the dendritic trees of pyramidal neurons (top left, triangles) in the two species are comparable in size, a target neuron (gray) pools input from a smaller region of visual space in ferret (top row) than in mouse (middle row). The synaptic receptive fields on the dendritic tree of the target neuron are distributed in a small (ferret, bottom middle) or large (mouse, bottom right) region in visual space. The receptive field spread is quantified by the standard deviation of the distribution of their centers, *σ*_*p*_ (5.3° for ferret and 26° for mouse, data reproduced from ref. ^(6,7)^). Example demonstrating lack of orientation clustering on a linear dendrite using mouse cortex receptive field spread over two simulated weeks. (**C**) Orientation difference between pairs of synapses as a function of distance in the model and data (data reproduced from ref. ^(6)^). (**D**) Correlations between pairs of synaptic inputs in the model and calcium signals of spontaneous synaptic activity in the mouse visual cortex (reproduced from ref. ^(3)^). (**E**) Receptive field overlap for pairs of synaptic inputs as a function of distance in the model and data (data reproduced from ref. ^(6)^).

In summary, a bigger spread of receptive field centers of dendritic synaptic inputs, as measured in the mouse cortex, produces a stable configuration of synapses lacking orientation clustering, but exhibiting receptive field overlap clustering. The anatomical argument suggests, that while synapses may not be clustered according to a particular stimulus feature in a particular sensory area^(13)^, synapses might still be clustered according to a different feature that is more relevant to the computations performed in that area. Quantitative similarities between synaptic clustering in both the developing and the adult mouse visual cortex strongly suggest that mature clusters are most likely formed during development through activity-dependent plasticity mechanisms that utilize developmental spontaneous activity patterns.

### Back-propagating action potentials establish global orientation clustering

We next asked whether the local organization of synaptic inputs achieved by our model also supports the emergence of global organization of synaptic inputs on the dendritic tree. To probe the interactions between the soma and synapses on different dendritic branches in a biologically realistic framework beyond the linear dendrites considered so far, we implemented our plasticity model on a morphologically realistic layer 2/3 pyramidal neuron (Allen Cell Type database, ID 508794889, Fig. 6A). We modeled a somatic signal that affects the dendrite in the form of backpropagating action potentials (bAPs) whenever the linearly summed input over all synapses exceeds a fixed threshold (Fig. 6A, inset bottom). These bAPs result in a calcium influx into the proximal and distal dendritic branches that attenuates with distance from the soma^(51)^ (Fig. 6B). As a result, the calcium signal at proximal synapses in our model neuron is dominated by somatic activity, while distal synapses are almost independent of the soma (Fig. 6C) in line with experiments^(52)^.

**Figure 6:**
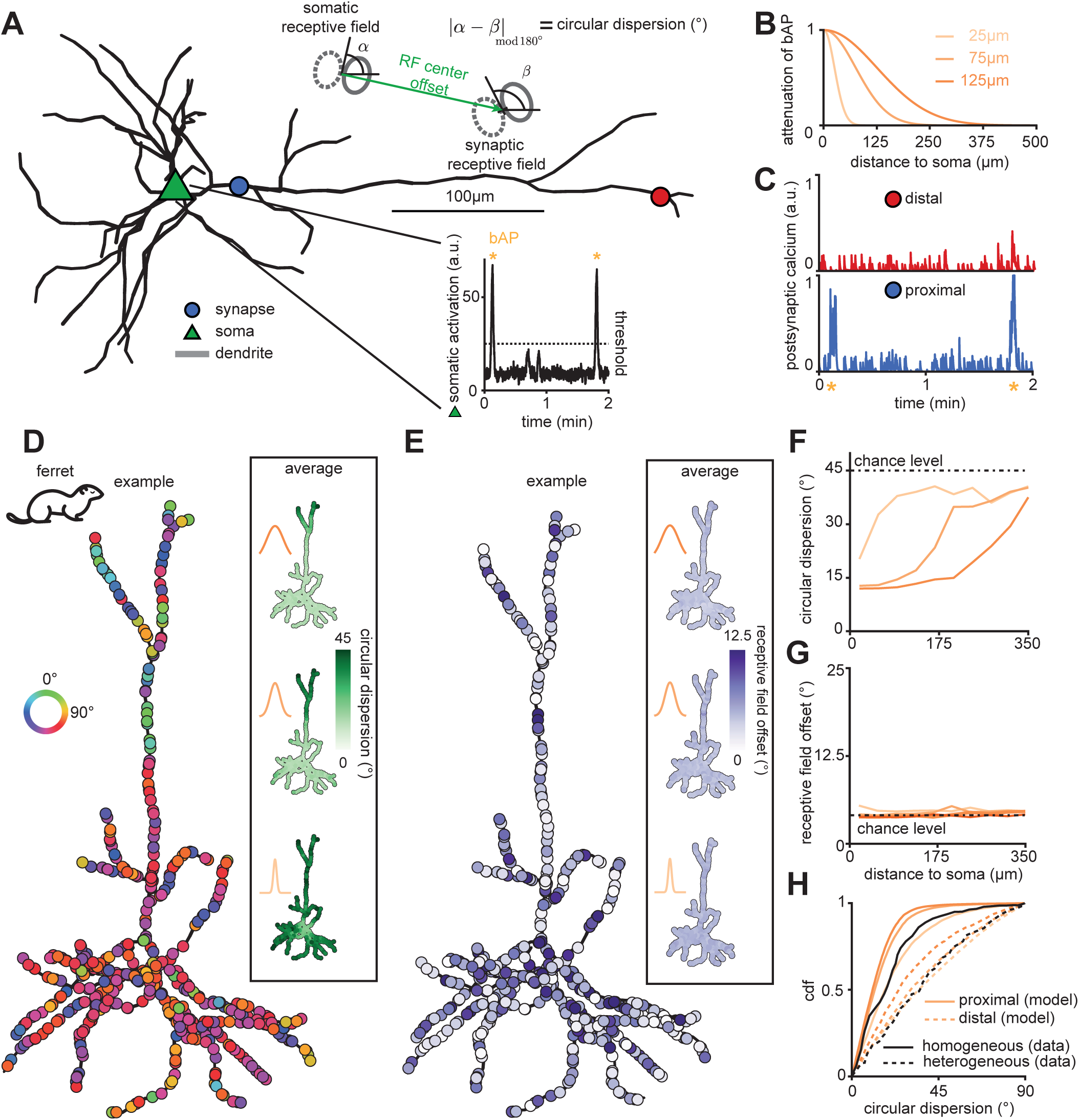
A backpropagating somatic signal drives heterogeneously and homogeneously clustered dendritic branches. (**A**) Illustration of a reconstructed pyramidal cell from layer 2/3 (Allen Cell Type database, ID 508794889). Triangle indicates soma, circles indicate synaptic sites. Inset top: Schematic to illustrate circular dispersion (see **D,F** and Methods) and receptive field offset (see **E,G**). Inset bottom: Sample trace of somatic activation under retinal wave stimulation after five simulated days. Stars indicate the initiation of a global somatic signal in the form of a bAP. (**B**) Attenuation of bAPs as a function of the path length along the dendrite for different attenuation factors. (**C**) Traces from the postsynaptic calcium level of a distal and a proximal synapse (indicated in **A**) for the same time period as in the inset of **A**. (**D**) Emergence of orientation clustering on the reconstructed pyramidal cell with synapses having a small receptive field center spread *σ*_*p*_ = 5.3° corresponding to ferret. Color indicates the orientation preference of the associated receptive field. Inset shows the circular dispersion averaged over 62 simulations for the three different attenuation factors from **B**. (**E**) Same as **D** but now color indicates the receptive field center offset. (**F**,**G**) Circular dispersion (**F**) and receptive field offset (**G**) between synapse and soma for different 1pa0th distances from the soma and different bAP attenuation factors. Same colors as in **B**. (**H**) Cumulative distribution function of circular dispersion for different bAP attenuation factors from **B** (data reproduced from ref. ^(5)^).

Using a small spread of receptive field centers in visual space corresponding to the ferret visual cortex (Fig. 5A), our plasticity model generates orientation clusters along the entire dendritic tree just like the linear dendrite (Fig. 6D, Supplementary Fig. 13). In addition, global organization also emerges. Synaptic inputs on proximal dendritic branches acquire orientation preferences similar to the soma due to the weak decay of bAP signaling, while inputs on distant dendrites have orientation preferences that are independent of the soma due to the strong bAP attenuation (Fig. 6D). Therefore, including bAPs homogenizes proximal dendritic branches to the orientation preference of the soma (Fig. 6D, inset). We characterized the degree of heterogeneity by computing the circular dispersion, i.e. the difference between the orientation preference of individual synapses and the soma (Fig. 6A, inset top and Methods). We find that bAP attenuation controls the extent of cluster heterogeneity along the dendritic tree (Fig. 6F). Consistent with experimental reports^(7)^, our model does not result in organization of the receptive field offsets, which are randomly distributed for synapses along the entire dendritic tree (Fig. 6E,G).

Different degrees of heterogeneity of synaptic orientation preference have been reported in the ferret visual cortex^(5)^, where experiments distinguished between *homogeneous* branches, with a mean circular dispersion below 15^°^, and *heterogeneous* branches, with a mean circular dispersion above 30^°^. We compared the cumulative distribution functions of the homogeneous and the heterogeneous branches to proximal (less than two times the attenuation constant) and distal (more than two times the attenuation constant) synapses in our model, respectively, and found good quantitative agreement (Fig. 6H). Thus, our results predict that bAP attenuation in different neurons may underlie branch orientation heterogeneity.

In summary, our model can produce local orientation clustering of synaptic inputs on a morphologically realistic model of an L2/3 pyramidal neuron, as well as global synaptic organization with synapses on proximal dendrites sharing similar orientation with the soma. We predict that an attenuating somatic signal is the main factor behind homogenizing proximal dendrites of the tree to the same orientation preference as the soma, while leaving distal synapses more heterogeneous and uncorrelated to the soma. This partial decoupling of dendrites from the soma has interesting implications for trial-to-trial variability and coding^(5)^, selectively enhancing computations through the generation of dendritic spikes^(53)^, or even predictively encoding specific features of the stimulus^(6)^.

### Emergence of visual topography of receptive field centers on mouse cortical dendrites

In addition to differences in local organization between mouse and ferret, the dendritic trees of the two species also exhibit a difference in global organization. In contrast to the lack of global organization of receptive field centers in the ferret^(7)^ (Fig. 6E,G), in pyramidal neurons of the mouse visual cortex, proximal vs. distal synapses respectively have a tendency to respond to more central vs. peripheral regions of visual space relative to the somatic receptive field center^(6)^. Additionally, peripheral synapses whose receptive fields are displaced along or close to the axis of the somatic receptive field, tend to share the same orientation preference as the soma^(6,54)^. To study whether a somatic signal like a bAP can induce such organization in our model of mouse pyramidal neurons, we considered a large spread of receptive field centers in visual space measured in mouse (Fig. 5A). We find that the bAP signal compels receptive fields of proximal synapses to be on average centered closer to the somatic receptive field compared to the receptive fields of distal synapses (Fig. 7C). This relationship is modulated by bAP attenuation, with weak attenuation increasing the number of synapses centered near the somatic receptive field (Fig. 7C,D). We refer to this global organization as a *dendritic map* of visual space, borrowing the term from the well-studied cortical maps^(55)^.

**Figure 7:**
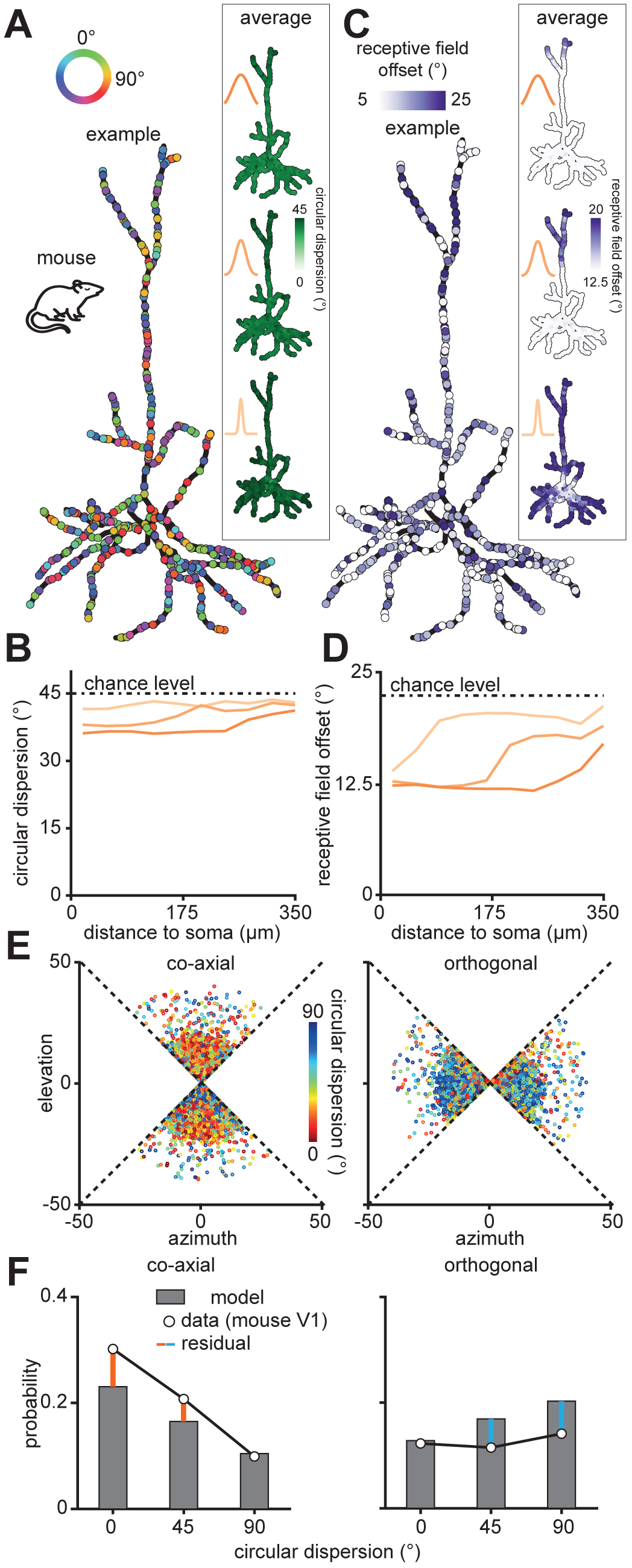
Backpropagating action potentials couple visual and dendritic space and establish a coarse dendritic map in the mouse. (**A**,**C**) Circular dispersion (**A**) and receptive field offset (**C**) on the reconstructed pyramidal cell with synapses having a large receptive field center spread *σ*_*p*_ = 26° corresponding to mouse. Colors and insets as in Fig. 6D and E. Data averaged over 80 simulations for the three different bAP attenuation factors. (**B**,**D**) Circular dispersion (**B**) and receptive field centers (**D**) for different bAP attenuation constants and path distances from the soma. Colors as in Fig. 6 F and G. (**E**) Receptive field centers in co-axial (left) and orthogonal (right) visual space colored according to their circular dispersion. (**F**) Histogram of circular dispersions (data reproduced from ref. ^(6)^).

As for the linear dendrite (Fig. 5B,E), synaptic inputs on the reconstructed pyramidal neuron exhibit no local orientation clustering (Fig. 7A) but robust overlap clustering (Supplementary Fig. 14). Consequently, bAPs in the model also fail to globally homogenize synaptic orientation preference on the dendritic tree and only introduce a bias to the overall distribution of orientation preferences as observed experimentally ^(6,14,15,49)^ (Fig. 7B). We hypothesized that homogenization might still occur in the mouse model but for receptive field overlap. Indeed, we find that while the absence of a bAP only generates local overlap clustering (Fig. 5C), including a bAP ensures that synapses close to the soma exhibit overlapping receptive fields with many other synapses close to the soma (Supplementary Fig. 14).

As a result of such globally homogeneous receptive field overlap, synapses with nearby receptive field centers in visual space have a similarly oriented receptive field to each other (Supplementary Fig. 15). We consequently observed that the distribution of receptive field orientations in different regions of visual space generated in the model has an interesting structure. In particular, synaptic receptive fields positioned along or close to the axis of the somatic receptive field, called the co-axial space (Supplementary Fig. 15), tend to have a similar orientation to the soma (Fig. 7E, left). Likewise, synapses in the region orthogonal to the co-axial space, called the orthogonal space (Supplementary Fig. 15), are more likely to be oriented orthogonally to the soma (Fig. 7E, right). Comparing the distributions of circular dispersions in the two regions of visual space to experiments^(6)^ produced good quantitative agreement, although large orientation differences were slightly overrepresented, while small orientation differences were slightly underrepresented (Fig. 7F). A possible explanation for this discrepancy might be a shift of the input distribution to a visual cortex pyramidal neuron after eye opening – when experiments were performed – in favor of inputs with small orientation differences^(43)^.

Taken together, our model provides a unifying framework for the emergence of local and global organization of synaptic inputs on the dendrites of cortical neurons in the mouse for different features than the ferret. In our framework, both the emergence of a retinotopic gradient along the dendritic tree, as well as the accumulation of co-aligned synapses in the co-axial portion of visual space, are explained through the homogenizing influence of bAPs. This has strong implications for the computational strategies of these neurons, for instance, the amplification of elongated edges prominent in natural stimuli, and how these neurons are integrated into local microcircuits.

## Discussion

Dendritic compartmentalization of synaptic inputs achieved by clustering has been postulated to enhance the computational capacity of neurons^(18)^, yet direct experimental evidence of clustering has only recently emerged. Despite functional synaptic clustering in the developing hippocampus^(1–3,8)^, no such fine-scale spatial organization was initially observed in the mature sensory cortex^(12–14,16)^. The emerging evidence now points to species-specificity of synaptic clustering for different stimulus features^(6,7,9,10)^. To reconcile these findings, we developed a unifying computational framework for the emergence of both local and global synaptic input organization on cortical dendrites. A single parameter, the cortical magnification factor of visual space, which is generally large for animals with large cortices and small for animals with small cortices, could explain local functional clustering for orientation in ferret vs. receptive field overlap and visual space in mouse cortex^(6,7,10)^. Interestingly, this magnification factor is also linked to the degree of input heterogeneity which can explain the population-level (columnar vs. salt-and-pepper) organization in the ferret vs. mouse visual cortex^(56)^, indicating that differences in the organization of dendritic synapses might result from the same universal developmental process modulated by evolutionary variations of cortex size. Our model also generates global organization of synaptic input feature selectivity mediated by an attenuating somatic signal. Global order is also achieved for different features in ferret and mouse cortex, effectively establishing soma-to-dendrite maps of orientation selectivity and visual topography, respectively.

### Generality of the minimal model

Our plasticity model was inspired by the push-pull mechanism of interacting neurotrophins operating in postnatal development^(8)^. We predict that early postnatal development is a particularly important time for the ubiquitous formation of synaptic clusters because the structural turnover of synapses occurs much more rapidly than in adulthood^(47)^. Thus, failure to correctly position synapses in development has severe and mostly irreversible consequences during later life^(57)^ and could play a role in neuropsychiatric disorders^(11)^. However, the minimal model derived from the neurotrophin model is sufficiently general to also be implemented in the adult, where the functional roles of MMP9 and calcium could equivalently be filled by postsynaptic depolarization, NMDA receptor activation, endocannabinoid or nitric oxide signaling or by more complex diffusible plasticity-related products^(58)^. Thus, our predictions regarding the organization of synapses are not contingent on a specific biophysical implementation of our minimal model as long as it has the three ingredients: distance- and correlation-dependent competition and structural plasticity.

### Origin of clustered synaptic input

The major source of excitatory inputs to pyramidal neurons in layer 2/3 neuron come from layer 4 neurons^(54)^, which likely obtain their orientation selectivity by combining On and Off center-surround receptive fields of thalamic feedforward inputs^(59)^. Our model also supports the concurrent emergence and clustering of orientation selectivity in the cortex, through the clustering of On and Off thalamocortical synapses (Supplementary Fig. 9). Also, pyramidal cells in the mouse and ferret visual cortex are already orientation selective at^(43)^ and before^(44,45)^ eye opening, providing another likely source of the synaptic Gabor receptive fields in our model. We find that a rapid increase in spine density during the second postnatal week^(60)^, which possibly indicates the establishment of dense, intralaminar and intercortical connections^(61)^, does not disrupt already established synaptic clusters (Supplementary Fig. 16).

### Functional role of clustered synapses

We propose three situations where synaptic input clustering may be beneficial for a neuron. (1) The transient, precise synchronization of even a small group of synapses can result in the nonlinear summation of synaptic activity^(62)^, enhancing a neuron’s computational capacity^(18)^ (however, see refs. ^(9,63)^). These nonlinearities can counteract location-dependent gradients of conductances across synapses, effectively establishing a synaptic democracy^(9,64)^. (2) Since synaptic transmission is highly variable^(65)^, multiple synapses encoding a similar signal might compensate for the potential failure of each individual synapse, thus effectively boosting the signal to noise ratio of synaptic transmission. (3) Furthermore, since the translation of proteins is localized to individual dendritic compartments where nearby synapses share available proteins, synaptic clustering is also beneficial from the perspective of resource-preservation^(66)^.

### Inhibitory synapses could form a backbone for excitatory clustering

Our model focused on the emergence of fine-scale organization of excitatory (glutamatergic) inputs, primarily due to lack of experimental data of the role of inhibitory (GABAergic) synapses on clustering. We speculate, however, that inhibitory synapses might shape the clustering of excitatory synapses on the dendritic tree. Indeed, while not being clustered themselves, GABAergic synapses can constrain the orientation preference of nearby excitatory clusters in our model (Supplementary Fig. 17). Since GABA might be excitatory early in postnatal development^(67,68)^, we considered two scenarios in the model with GABA switching from excitatory to inhibitory, or being inhibitory the entire time. The resulting excitatory clusters are tuned to the same orientation relative to nearby inhibitory synapses in the former case or tend to prefer the orthogonal orientation in the latter case (Supplementary Fig. 17). Thus, we predict that GABAergic synapses might be co-clustered with excitatory synapses. Provided that GABAergic synapses fire in synchrony with glutamatergic synapses, this co-clustering could allow them to dynamically switch a given cluster ON or OFF^(69)^. Additionally, since inhibitory synapses can cancel the effect of backpropagating action potentials^(70)^, we expect that inhibitory synapses synchronized with somatic activation would be able to protect a cluster from the homogenizing effect of backpropagating somatic signals^(71,72)^.

In summary, our model unifies several experimental studies from the last decade on the emergence of functional synaptic organization for different features, at different scales and in different species. It makes the surprising prediction that synaptic clustering observed during development is qualitatively similar to clustering in the adult suggesting that similar mechanisms are in operation. Therefore, a single framework can explain how circuits can wire up with sub-cellular precision, with paramount implications on the computational properties of cortical neurons and networks.

## Acknowledgments

We thank all members of the Gjorgjieva lab, M. Kaschube and G. Laurent for comments and discussions, and M. Jüngling, J. Letzkus, C. Lohmann, C. Miehl, M. Silies and M. E. Wosniack for feedback on the manuscript. This work was supported by the Max Planck Society, SMART START training program in computational neuroscience (to JK), the European Research Council (StG 804824 to JG) and the Behrens-Weise Foundation (to JG).

## Author Contributions

JK and JG designed the research. JK analyzed the model and performed model simulations. JK and JG prepared the figures and wrote the manuscript.

## Methods

### Neurotrophin model

We base our neurotrophin plasticity model on interactions between signaling molecules shown to drive the emergence of synaptic clustering during development^(1–3,8)^: brain derived neurotrophic factor, BDNF (*B*), its immature form, proBDNF (*P*), the cleaving protease matrix metallopeptidase 9, MMP9 (*M*), and postsynaptic calcium (*Y*). *W*_*k*_ is the synaptic efficacy of a synapse with hard bounds at zero and one. For a pair of synapses *k* and *l* separated by *d*_*kl*_ along the branch, we define the proximity variables 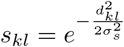, where (*σ*_*s*_ determines the spatial postsynaptic calcium spread constant.

Since the mechanism that produces synaptic clusters is activity-dependent, we model presynaptic and postsynaptic accumulators of synaptic activity. We model the presynaptic accumulator MMP9 as a synapse-specific leaky accumulator^(25)^ with dynamics

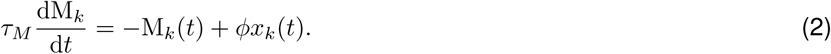

Here, 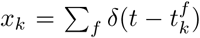 is the input even train to the *k*-th synapse with events at times 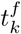, and *ϕ* is an MMP9 efficiency constant that determines how efficiently MMP9 converts proBDNF into BDNF. The postsynaptic accumulator is modeled by the local calcium variable *Y* that integrates activity from nearby synapses^(26)^, weighted by their efficacies *W*_*l*_ and the distance-dependent factor *s*_*kl*_,

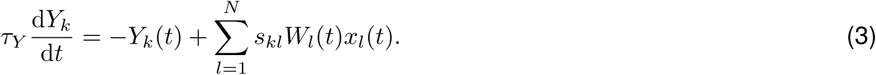

Following experimental data^(8,32)^, we couple extracellular proBDNF and BDNF to postsynaptic calcium and let MMP9 convert proBDNF to an equal amount of BDNF,

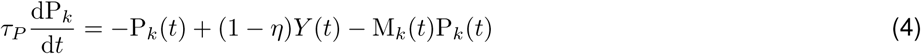

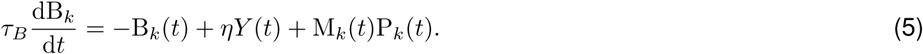

Here, the scaling factors (1 − *η*) and *η* ensure that without MMP9 the ratio of BDNF to proBDNF remains close to the constitutive ratio *η*^(73)^. According to the Yin-Yang hypothesis of neurotrophin action^(23)^, binding of BDNF to its receptor TrkB leads to synaptic potentiation, while binding of proBDNF to p75^NTR^ leads to depression. We assume that the binding affinities and corresponding magnitudes of induced plasticity are balanced, so that the synaptic efficacy can be written as the difference between BDNF and proBDNF,

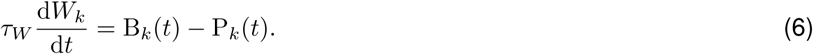

*τ* always denotes the time constant for the variable in the corresponding subscript. The values for all parameters used in our simulations are listed in Table 1.

### Minimal model

Assuming a tight coupling of proBDNF and BDNF to the postsynaptic calcium^(32)^, we make a steady-state approximation of *P*_*k*_ and *B*_*k*_ in equations (4) and (5), insert these expressions into equation (6) for *W*_*k*_ and then linearize the resulting function around *M*_*k*_ = 0 to obtain a minimal model (see Supplementary Note 1 for details). While we use upper case letters for the variables in the full neurotrophin model, we use lower case letters for the minimal model. The minimal model consists of a synapse-specific presynaptic accumulator *v*_*k*_,

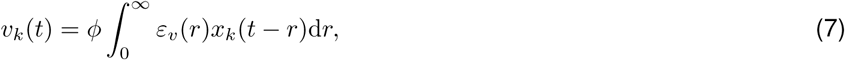

where 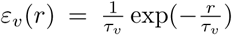 for *r* ≥ 0 denotes the time course of an excitatory presynaptic event and a postsynaptic accumulator *u*_*k*_ that averages over nearby synapses in a weighted and distance-dependent manner,

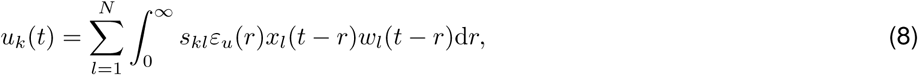

where 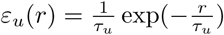 *r* ≥ 0 denotes the time course of an excitatory postsynaptic event. The efficacy equation (6) turns into a Hebbian equation that directly combines the pre- and postsynaptic accumulator with an additional offset constant *ρ*,

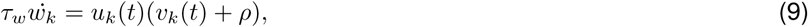

with 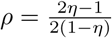 and 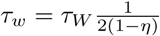. This model is minimal in the sense that it cannot be reduced further without losing either the dependence on correlation through the link to the BTDP rule, or the dependence on distance.

### Steady-state analysis of the minimal model

Combining the equations for the accumulators and the efficacy dynamics (7)-(9), taking the expected value over neurons (denoted by ⟨·⟩) and over time (denoted by 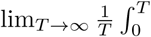 ·d*t*) we can write the expected change in synaptic efficacy as (see Supplementary Note 2 for full derivation)

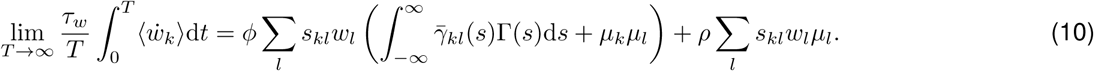

Here, 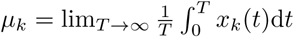 is the mean firing rate of the *k*-th input 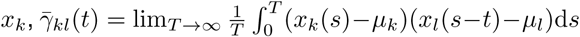 is the covariance between inputs *k* and *l* at lag *t* and the kernel 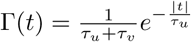.

When only one input is activated with burst events of duration *x*_dur_ and rate *μ*, we simplify equation (10) to write the change in synaptic efficacy for this input as *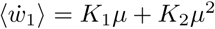*, while the change in efficacy for a second inactive input at a distance *d*_12_ is *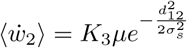*, where *K*_1_, *K*_2_ and *K*_3_ are constants (see Supplementary Note 3 for details), as shown in Figure 1D.

For the completely homogeneous case with identical inputs on a linear dendrite with density *v*, equal efficacies *w*_*k*_ = *w* and rates *μ*_*k*_ = *μ* for all *k*, and identical correlation *c*_*kl*_ = *c* for all pairs *k* ≠ *l*, equation (10) can be used to derive the critical amount of correlation *c*\* at which the system switches from the depression-dominated into the potentiation-dominated regime, *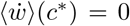*, as 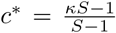 where 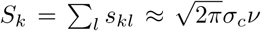, which is the same for all inputs *k*, thus *S* = *S*_*k*_, see Supplementary Note 4. *S* can be thought of as a measure of the total amount of activity in an area around a given synapse and 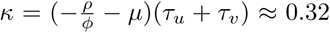 is a constant. Note that for high densities *v*, the critical value *c*\* quickly approaches *k* and is bounded above by it. The expression for *c*\* provides the dashed line in Figure 3B, while the other contour lines stem directly from equation (10) and the approximation 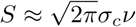 (see Supplementary Note 4).

To analytically approximate the change in synaptic efficacy for synaptic pairs with heterogeneous correlations and initial efficacy (Fig. 3E, right), we use the steady-state equation (10) where we insert the local average correlation and the local average initial efficacy to obtain a linear expression 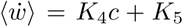, where *K*_4_ and *K*_5_ are again constants (see Supplementary Note 4).

### Simulations

For the simulations in Figure 1B,D we distribute two synapses at varying distances (Δ*d* =0 μm to 15 μm) on a linear dendrite of length *L* =150 μm with periodic boundary conditions. Only one input is stimulated with a train of 50ms-long burst events whose rate varies from 1 min^−1^ to 20 min^−1^. We fix the synaptic efficacy to the initial value *W*(*t*) = *W*_0_ and compute the expected change in the efficacy of synapse i in equation (6) as the temporal average of B_*k*_(*t*) − P_*k*_(*t*) over the 40min duration of the simulation. For the simulations in Figure 2B-D we simulate only one synapse and provide either one (panel b and c) or ten (panel d) pairings of a pre- (*t*_*pre*_) and a postsynaptic (*t*_*pre*_) burst event at a temporal offset of Δ*T* = *t*_*post*_ − *t*_*pre*_. Each burst event has a duration of one second and contains ten smaller 10ms-long events. The postsynaptic burst contributes an additive term *I*(*t*) with amplitude *B*_*amp*_ to the postsynaptic calcium, 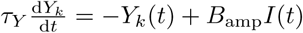.

For the simulations in Figure 3B,C, we distribute *N* = ⌊*Lv*⌋ synapses on a branch of length *L*, where ⌊*x*⌋ is the largest integer smaller or equal to *x* and where we choose *L* sufficiently long (30 μm) to avoid confounds from periodic boundary conditions. We vary the density *v* between 0.05 μm^−1^ and 0.75 μm^−1^ to capture random and systematic fluctuations in local synapse density^(76)^. We generate 12 minutes (4 hours in Fig. 3C) of correlated Poisson event trains^(77)^, homogeneous across pairs and with a fixed firing rate of 15min^−1^, and compute the expected value of the change in synaptic efficacy via equation (9) over time. We also test the case of heterogeneous correlations across different pairs (Fig. 3D,E). For this, we sample pairwise correlations from a beta distribution *B*(*U*, 10) where the first shape parameter *U* is itself a uniform random variable that takes values from the interval [1, 4] (Fig. 3D) or [0, 100] (Fig. 3E) to obtain local correlations with small or large magnitude. We use these values to form the upper triangle of the matrix, and copy them into the lower triangle matrix to obtain a symmetric matrix from which 24 minutes (Fig. 3D) or 12 minutes^(3)^ (Fig. 3E) of Poisson events were produced^(77)^. The initial synaptic efficacy in Figure 3B-D is 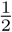; in Figure 3E the initial efficacy is uniformly sampled in a broader interval [0.25, 0.75].

In Figure 4 and Figure 5, we simulate a branch with length L of 150 µm over 15 days. For the simulations in Figure 6 and Figure 7, we test a morphologically realistic dendrite model by using a reconstructed pyramidal cell from layer 2/3 of the mouse visual cortex (Allen Cell Type database, ID 508794889) which we resample into equally sized segments of 5 µm using the TREES toolbox^(78)^. We use this dendritic tree for both types of simulations, ferret and mouse, since to our knowledge no morphological reconstruction of a pyramidal layer 2/3 neuron from the ferret visual cortex is openly available. We do not expect this to influence our results, since using the dendrite of a layer 6 pyramidal neuron from the ferret visual cortex (NeuroMorpho.Org ID NMO_105857) still produces local and global clustering (Supplementary Fig. 18). Because of the increased computational complexity, the duration for the simulations with the morphologically realistic dendrite model is 5 days, which is sufficient for most of the synapses to reach a stable state (Supplementary Fig. 19).

To mimic the density of excitatory synapses before eye opening in the visual cortex^(76)^, in Figures 4 to 7 we fix the average density of synaptic inputs to *ν* =0.2 µm^−1^. The newly generated efficacy of a synapse through structural plasticity is the same as the initial efficacy at the onset of the simulation, i.e. 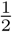. Similarly, for both initial and newly generated synapses, the orientation of the receptive field is drawn from a uniform distribution over the inverval [0°, 360°]. Synaptic receptive field position is known to be spread out in a small (for ferret^(7)^) or large (for mouse^(6,54)^) neighborhood around the somatic receptive field center. We define the receptive field center spread *σ*_*p*_ to incorporate these differences and estimate *σ*_*p*_ from experimental studies in both species^(6,7)^ as the standard deviation of a Gaussian (Fig. 6A). The parameter *σ*_*p*_ is inversely related to the cortical magnification factor of visual space, since a larger cortical magnification results in a smaller spread of receptive field centers of the synaptic inputs for any given neuron^(6,7,56)^ (Fig. 6A). Consequently, an initial or newly generated synaptic receptive field center is drawn from a two-dimensional symmetric Gaussian with standard deviation *σ*_*p*_ and truncated to a circle of radius 50° to ensure that receptive fields fall within a region of visual space where they are modulated by the retinal waves.

### Statistics

All correlations are computed as Pearson correlation coefficients, i.e. for two random variables *X* and *Y* we compute 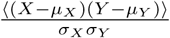, where *µ* and *σ* denote the mean and the standard deviation. In Figure 3B, for each pair of correlation and synaptic density we compute the average change in synaptic efficacy over all synapses in 50 simulations. In Figure 3D,E the term ‘local correlation’ refers to zero-lag correlations between input trains, averaged in a neighborhood of 6 µm. Each point is a synapse from one of 30 synapses per simulation in 10,000 simulations.

To compute the correlations in Figure 4B,G and Figure 5D, we first apply a boxcar filter of length 3 s to generate signals consistent with the slow calcium dynamics in experimental imaging studies^(14)^. The experiments from ref. ^(3)^ reproduced in Figure 3E and Figure 5D reports *coactivity* – the fraction of events at a given synapse that occurs in concert with events at nearby synapses. Coactivity is closely related to the Pearson correlation coefficient when it is only applied to pairs of synapses (see Supplementary Note 5).

We compute the receptive field overlap in Figure 5C as the spatial receptive field correlation^(6)^, i.e. the pixel-wise Pearson correlation coefficient between Gabor filters associated with pairs of synapses. The term *orientation difference*^(6)^ denotes the absolute difference in orientation between receptive fields of pairs of synapses modulo 180°,

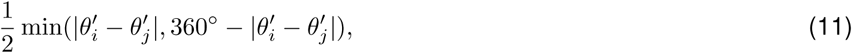

where 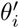 is defined as 2mod(*θ*_*i*_, 180°). The term *circular dispersion* denotes the orientation difference between the receptive field of a given synapse and the soma, where we compute the orientation preference of the soma as the circular average of all synaptic preferences^(5)^, 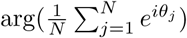. Analogously, the term *receptive field offset* denotes the Euclidean distance between the center of a given synaptic receptive field and the somatic receptive field center, defined as the average location of all synaptic receptive fields. The average circular dispersion and average receptive field offset in Figures 6 and 7 are computed as the average over synapses within 5 µm dendritic segments. The *co-axial* space^(6)^ (Fig. 7E,F) is defined as the portion of space up to 45° on either side of the axis extending along the average orientation of all synaptic receptive fields. Conversely, the *orthogonal space* is the remaining visual space that is not co-axial.

We extract all displayed experimental data from the appropriately referenced publications using open source software^(79)^.

### Retinal wave generation

To generate retinal waves with realistic spatio-temporal properties, we simulate six hours of retinal waves from a published computational model with the parameter setting for mice (P0-P13)^(41)^ and loop the waves over the entire duration of the simulation. For consistency, we use the mouse retinal wave parameter settings for all simulations, since the parameter settings for ferret stem from much younger animals (P2-P4)^(41)^. Using the parameters appropriate for ferret still generates synaptic clusters, albeit with slightly larger orientation differences between nearby synapses (Supplementary Fig. 20). As a control we also generate white noise input by sampling each pixel independently from a normal distribution and applying a spatial Gaussian filter with a 2° standard deviation. Subsequently, we use each frame of the retinal wave or the white noise movie as input to a linear-nonlinear Poisson model of event generation.

### Linear-nonlinear model

The linear filter consists of a Gabor linear filter H, constructed as the difference of two Gaussian distributions with semi-minor and semi-major axis lengths *σ*_*min*_ and *σ*_*maj*_. For consistency, we use the same lengths of the semi-major and semi-minor axes of the Gaussian for both mouse and ferret (Fig. 4A and Table 1). This is supported by experimental evidence^(6,7)^ which shows that the receptive fields of synaptic inputs of mouse and ferret do not differ by more than a factor of 1.5. Each individual Gabor filter is rotated according to its orientation between 0° and 360*°, θ*, and has a shifted receptive field center with azimuth and elevation between −50° and 50°. The linearly filtered stimulus is passed through an exponential nonlinearity, *a* exp(*bH*), that produces an instantaneous firing rate from which we generate a Poisson input train with individual 50ms-long events.

### Structural plasticity

To model synaptic turnover, we implement a structural plasticity rule inspired by ref. ^(47)^ where each synapse whose efficacy falls below a fixed threshold *W*_thr_ is removed and replaced by a new synapse with a random position on the branch and a randomly oriented Gabor receptive field (Fig. 4D). This novel synapse could potentially come from a pool of *silent synapses*, a type of synapse that lacks AMPA receptors but can become unsilenced through activity-dependent mechanisms^(80)^. While we maintain a constant density of synapses throughout our simulations, a rapid increase in synaptic density like the one observed during development in the somatosensory cortex^(61)^ does not interfere with the formation of synaptic clustering (Supplementary Fig. 16).

### Backpropagating somatic signal

To investigate the emergence of global order of synaptic inputs on an entire dendritic tree, we add a somatic accumulator which can produce backpropagating action potentials which attenuate with distance from the soma. Our somatic accumulator sums linearly over all postsynaptic accumulators, weighing them by their respective synaptic efficacy^(18,81)^

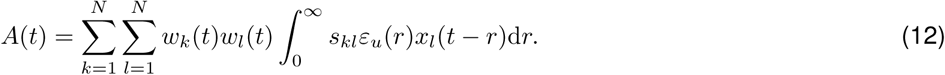

If the somatic accumulator crosses a threshold, *A*_th_, there is a 25% probability that a backpropagating action potential *B*(*t*) ∈ {0, 1} is generated, which affects the postsynaptic accumulators of all synapses with attenuating effect over distance^(51,52)^

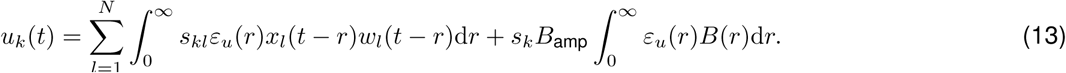

Here, *B*_amp_ is the unattenuated strength of the bAP and 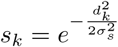 is the attenuation factor of synapse *k* that depends on the distance to the soma d_*k*_ (Fig. 6B).

